# A Preventive Effect of Exercise in a Mouse Model of Essential Tremor

**DOI:** 10.64898/2026.11.02.763708

**Authors:** Sofia Constantopoulos, Andreas Bergdahl, Richard Courtemanche

## Abstract

The effects of exercise have not been widely studied for potential benefits in essential tremor (ET), while this has been growing for other neurodegenerative disorders. This study examines the effects of two types of exercise [high-intensity interval training (HIIT) and resistance training (RT)] on mice that were subjected to harmaline injections to trigger tremor. We employed a repeated-measures design with three groups: a HIIT group, a RT group, and a sedentary (SED) control group. Each group received an injection before and after the intervention. Tremor kinetics (power within the 8-14 and 14-29 Hz bands) were assessed using accelerometry. Tremor kinematics were also assessed using high-speed video analysis, for validation. As potential signalling molecules from exercise, we also tested for blood lactate and BDNF in each group. Compared to pre-intervention measurements, we found a significant decrease in power within the 8-14 Hz band combined with an increase in the 14-29 Hz band, for the HIIT group, but not in the RT or SED groups. We also found a change in BDNF concentration for the HIIT group. These results suggest that the HIIT protocol may have helped the animal to shift tremor frequency to higher bands, thereby offering functional resistance to harmaline-induced tremor. This study supports high intensity aerobic physical exercise as a potential therapeutic/preventive intervention for ET, with translational implications for human studies.

**NEW & NOTEWORTHY:** We show that high-intensity interval training (HIIT) can help modulate tremor effects in the harmaline mouse model of essential tremor (ET), by shifting the dominant frequency band from lower to higher values. This was specific to HIIT, in contrast to resistance training. We also report that this could be partly due to an increase in exercise-induced BDNF, paving the way for the use of whole-body intense exercise to help minimize tremor in ET patients.

## INTRODUCTION

Essential tremor (ET) is the most common movement disorder characterized by an involuntary, rhythmic and oscillatory kinetic tremor (1). It affects up to 1% of the population, with prevalence increasing with age, particularly in individuals aged 65 and older (2), but also presents in children, with a mean onset age around 9.7 yrs (3). Animal models have been useful for deciphering the pathophysiology of ET; one such model, the harmaline-induced tremor model, has helped in understanding circuit-level reverberatory mechanisms (4, 5). Indeed, harmaline induces a transient, kinetic tremor by entraining cerebellar circuitry, notably affecting the inferior olive (IO), cerebellar cortex, and deep cerebellar nuclei (DCN) (6). In this model, the IO rhythmically entrains the sensorimotor networks, which allows for the investigation of circuit-level mechanisms (7), including the entrainment of thalamic oscillations (8). Despite the widespread use of this model, the exact role of the IO in ET remains debated, particularly as cerebellar overactivity, rather than IO oscillations, may drive the tremor (9, 10). Nonetheless, its attractiveness resides in how it is amenable to physiological enquiry, and the capacity to design time-constrained experimentation.

While pharmacological and surgical interventions for ET exist, they often carry significant side effects and are not universally effective (3). Exercise is increasingly seen as a way to improve overall brain health, making the brain more resilient to disease, prevent loss of function with age, and maintain multiple sensorimotor and cognitive abilities through the years (11–13). Physical activity may offer neuroprotective benefits in Parkinson’s disease and other neurodegenerative disorders: physical activity improves neuroplasticity and slows down disease progression (14–17), by modulating neuronal circuits, increasing neurotrophic factors, and decreasing neuro-inflammation (17, 18). In ET, the effect of exercise on tremor has been less explored, though there could be benefits, as evidenced by improvements in grip strength and other aspects of motor function in rodent models (19). For general health benefits, high-intensity interval training (HIIT) has emerged as an effective exercise protocol, providing multiple cardiovascular benefits and potential neuroprotective advantages (16, 20–23). In addition to entraining the sensorimotor circuits through coordinated movement control, HIIT could potentially enhance neuroprotection via its influence on hormones and circulatory factors (24, 25), also known as exerkines (26–28), including brain-derived neurotrophic factor (BDNF), which could potentially offer a protective effect against tremor (23). Recent research indicates that HIIT may improve motor performance and brain plasticity in Parkinson’s disease models, by acting as a countereffect to disease progression (29). Indeed, exercise as an intervention could prove optimal in setting a neural environment that could foster optimized patterns of neural activity (16). We hypothesize that similar preventive benefits may extend to ET, as both conditions dysregulate the sensorimotor networks, including through cerebello-cortical circuits, into excessive reverberation.

The specific aim of this study was to assess the preventive effects of a HIIT exercise program on tremor in a harmaline-induced mouse model of ET, in comparison with a resistance training protocol and a sedentary group. We hypothesized that the HIIT exercise protocol, through mechanisms favoring neuroplasticity, would attenuate tremor severity under a harmaline challenge, so circuits can freely function. By exploring exercise as a preventive intervention, we are targeting the capacity of controlled, titrated physical activity to preserve the voluntary control of neural circuits, via specific movement activation and exercise-related metabolic effects, to counterbalance the progression of neurodegeneration. This can further provide insight into mechanisms benefiting an overall intervention strategy in human ET patients.

## MATERIALS AND METHODS

This study was structured to test the effect of different types of training modalities relative to a harmaline challenge, before and after an exercise regimen for 2 weeks. We aimed to assess how a controlled exercise protocol could influence the tremor response to harmaline, using accelerometry. All methods adhered the guidelines of Concordia University Animal Research Ethics Committee, compliant with the regulations from the Canadian Council on Animal Care.

### Animals

This study involved 21 adult C57Bl/6 mice, with males (n = 2) and females (n = 19). The mice were aged 6-15 months, a range covering the adult onset to older period of ET (30). The mice weighed an average of 32.9 g pre-intervention, and 32.28 g post-intervention. They were housed in groups of 2-4 animals in 12h/12h lighting conditions and were trained and tested during the light phase. Mice were fed regular rodent chow and given standard cage enrichment. The mice were divided in 3 groups: a HIIT group (n=7), an anaerobic exercise (resistance training) group (RT, n=7) and a sedentary control group (SED, n=7).

### Study Design

To study the effects of the interventions on tremor, we followed a repeated-measures design with the 3 groups of mice. The relative timeline is detailed in Figure 1. Harmaline injections were given to each mouse before and after the 2 weeks of intervention, providing a comparable response to the harmaline challenge. Tremor was thus measured at 2 time points, after the pre- intervention harmaline injection, and after the post-intervention harmaline injection.

**Fig. 1.**
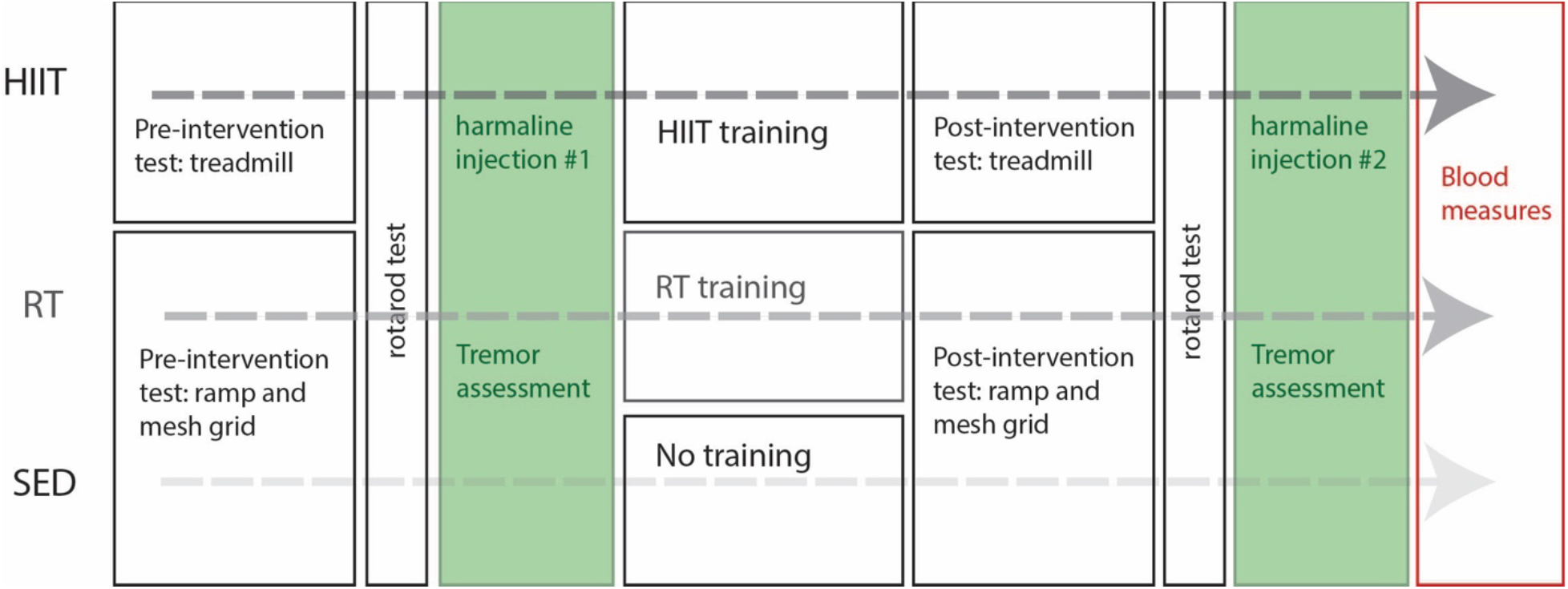
The general protocol followed for each group. For the HIIT group, mice ran on the treadmill to screen if they were eligible for aerobic exercise; if qualified, they performed an exhaustion test, were tested on the rotarod. Tremor assessment was performed in response to the harmaline injection, before and after HIIT training and post-intervention testing. A similar schedule was followed for the RT and SED groups, with appropriate RT-like testing, harmaline injection around the optimized intervention (RT), or nointervention (SED).

### Pre-intervention testing

Before beginning their assigned protocols, all mice underwent a pre-screening and pre-testing session to determine group allocation. The pre-screening process included a treadmill run to assess each mouse’s eligibility for the HIIT group, also serving as a first day of acclimation for these mice. The pre-testing process was comprised of two sessions performed on separate days. On the first day, the mice ran on the treadmill at a speed of 8 m/min during a 10-min warm-up. The speed was then increased to 15 m/min for 15 min and the session concluded with a 10-min cooldown at 8 m/min. On the second day, an exhaustion test was performed: this session consisted of the same as the first day 10-min warmup, five sets of 1-min intervals at 17 m/min separated by 2-min periods at slower pace (12 m/min), and a 10-min cool down. If any mouse failed to match the pace (touching the brush at the back of the corridor on the treadmill’s belt) even with tactile motivation (a small shake of the brush) for 10 s or more, they were considered unsuitable for inclusion in the HIIT group. These mice were then included in the RT or SED groups, who underwent circuit training as a pre-testing phase. The SED testing was to put in context the passage of time in the overall protocol.

To summarily assess the basic posture and movement capacity of the mice, we observed their movement and locomotion when walking in their cages; in addition, we placed them on a rotarod to assess their motor coordination and balance (31), at three different speeds (4, 8, and 12 RPMs), and measured the time they could stay on the rotating axis.

### Exercise Interventions

Overall, we had two intervention groups and one control group, and the exercise protocols follow the developing guidelines for animal models (32).

#### High-intensity interval training – HIIT

This group received a high-intensity aerobic exercise intervention, where mice underwent a 14-day protocol to increase cardiovascular and muscular fitness, with 3 sessions per week on a speed-controlled rodent treadmill (33). Each session lasted 25 min, starting with a 5-min warm-up (< 50% maximum running speed) followed by 5 sets of 1 min at high intensity (> 75% of maximum running speed), separated by 2-min active recovery periods, all finalized with a 5-min cool-down (34).

#### Resistance training – RT

This group received a muscle-strength targeted exercise regimen consisting of circuit-like training, with each session comprised of (1) hanging on a mesh grid (3 trials of minimum 60 s up to 180 s separated by 15 s of rest), and (2) climbing up a 90° ramp, (8 trials, unweighted, without any rest between the trials). One acclimation session to the apparatus was provided. After each session, a trial was added to increase the volume. Our initial protocol consisted of inclined ramp weight-pulling using homemade vests, to attach variable weights; however, our mice never acclimated in wearing the vest for multiple days, affecting their food and water consumption. This circuit-like training was to provide consistent muscle work with different exercises, using repetitions to increase the workload, all within a short time.

#### No exercise – SED group

This group was constituted as a comparative control for a lower level of activity. SED mice were kept in their home cage with casual handling for the same duration as the other groups (2 weeks). They also underwent the same pre- and post-testing as the RT group but did not perform either of the HIIT or RT protocols.

### Post-intervention testing

Before the second harmaline administration was given, all mice underwent post-intervention group-specific exercise testing, along with the rotarod test, to compare with pre-intervention values. Expectations were that the exercise capacity would be improved after the intervention.

### Harmaline injections

To initiate tremor, a solution of harmaline was injected intra-peritoneally [20 mg/kg of harmaline hydrochloride (Cedarlane Laboratories, Burlington ON, CAN)], as per previous approaches (6). As reported and at the right dosages, this drug produces body tremors in mice at 11-14 Hz, evident in the forelimbs, lasting 1-2 hrs and peaking at 30 min (35, 36). We expected to measure the tremor between the 25^th^ and 35^th^ min post-injection, but in our hands, an apparent tremor began faster, at the 5-min mark post-injection and the tremor was measured for a duration of up to 35-40 min post-injection. The peak was observed between the 15-20 min mark. Once the tremor visibly subdued, mice were put back in a transition cage, and then back to their group home cage after a maximum of 30 mins.

### Measurement of tremor

At the time of pre- or post-intervention tremor challenge, each mouse was placed in a specially designed setup for measuring tremor kinetics at the whole-body level. This enclosure was safe for the mice and they could move around freely, though mainly stayed relatively in place for many seconds. Tremor was measured for each mouse, using accelerometry (kinetics) and for certain validation experiments, video was analyzed (kinematics).

### Kinetics

A standard mouse plastic cage bottom was suspended by two elastics to a square wooden stand, similar to done previously (37). The suspended cage formed an enclosure free of ground reaction forces, permitting to measure the animal’s vibrations with tri-axial accelerometry. An iPhone 6 (Apple, Cupertino, CA, USA) was velcroed to the plastic cage’s underside, and its accelerometry measures were accessed with an application [Tremor 12, (38)]. This open-source mobile application allowed for measurements of rotation, acceleration, and gravity in all 3 axes (x, y, z) and timestamps were sampled at a 100-Hz frequency.

### Kinematics

During tremor assessment, digital video [recorded with a GoPro Hero8 camera (San Mateo, CA USA), with a 4K resolution at 60 frames/s] was taken of each mouse (placed in front, centered on the whole cage, for a constant field of view) to provide information as to the body configuration and tremor kinematics. For multiple sessions, video was visually inspected and quantified. Quick, brief analyses consisted in the frame-by-frame labelling of body landmarks for prospective time periods, with simple body marker identification and analysis in MATLAB (R2021a, MathWorks, Natick, MA, USA). For more extensive analyses, we used an open-source deep-learning Python-based analysis environment for markerless pose estimation called DeepLabCut (39). This approach requires training of a model, which we performed on an Alienware (Miami, FL, USA) Aurora R15 computer with a NVidia GeForce RTX 4080 GPU. In DeepLabCut, our typical labelling of 50 frames trained a model to track the nose, the ears, top of the back, the left and right flanks, the tail base and tip, and the forepaws of the mice. This larger-scale approach was used on a few videos to complement the primary measurement of tremor using the analysis of accelerometry, which was done on all animals.

### Physiological measurements: blood lactate and BDNF

In order to address the potential effect of exercise training on the brain, the link with brain plasticity was explored through the measurement of lactate produced after exercise (HIIT and RT groups) or daily activity (SED group), as well as the measurement of circulating BDNF, a growth factor that has been linked to exercise (15, 16).

A tail bleed was performed on each mouse to collect a drop of blood which was then transferred onto a lactate strip and inserted into the lactate meter (Lactate Plus Meter and test strips, Nova Biomedical Canada; Mississauga, ON, CAN). For certain mice, lactate measurements were performed at the time of euthanasia, with blood coming from the cardiac punctures.

For each mouse, at the time of euthanasia, cardiac punctures were performed to collect blood for measurement of circulating BDNF. Approximately 80-1000 μl of whole blood was collected from each mouse. These samples were transferred into Eppendorf tubes and allowed to rest at room temperature for 30 mins, followed by 30 mins at 4°C. The blood was centrifuged to separate the blood cells from the serum at 2000 G for 10 mins. The serum was carefully collected from the top of the tube and centrifuged again at 2000 G for 10 mins to remove any remaining cells. The serum was kept at -80°C for later analysis (40). Upon having collected all blood samples from the mice, quantification of the BDNF concentration in the serum was assayed using a Biosensis mature BDNF Rapid ELISA Kit (human-mouse-rat, VWR Intl, Mississauga, ON, CAN). Horseradish peroxidase was employed as the antibody for detection. A serial dilution of standards was performed, ranging from 125 pg/ml to 1.95 pg/ml, to generate a standard curve. Quality control procedures were implemented to ensure assay accuracy and reliability, and for all but one mouse, multiple samples (most 2, some 3) were averaged.

### Data analysis and statistics

Most statistics were processed using SPSS (IBM, Armonk, NY, USA) using conventional tests. We focused on the differences between the three groups. For exercise and sensorimotor performance, the pre-intervention and post-intervention data (physical testing, rotarod testing) were submitted to a t-test for changes between pre-intervention and post-intervention, for each group. Accelerometry and kinematics required further process, described below.

For the tremor quantification, the raw measurements from the TREMOR12 software were exported to MATLAB (MathWorks, Natick, MA, USA). Visual inspection was performed on all the records. Using the *spectrogram* function, the spectrum of signal power vs. frequency was produced, for each recorded file [multiple consecutive 1-s windows with 50% overlap], with a focus on the acceleration in Z (ACCZ) and rotation in X (ROTX) variables; these showed the most distinctive tremor profiles during the time the animal was in the measurement apparatus. ACCZ represented the outcome of the force acting perpendicularly on the cage bottom, so perpendicularly to the accelerometer. ROTX showed responsivity to the force acting to rotate the cage along the axis of the two rubber bands. Specific frequency boundaries (8-14 & 14-29 Hz) appeared best suited to quantify the tremor for the mice in our apparatus. An example for each variable is shown on Figure 2. Both spectrograms (ACCZ and ROTX) provide evidence for the use of the specific boundaries at 8-14 Hz and 14-29 Hz, as they capture most of the rhythmicity components of the signal.

**Fig. 2.**
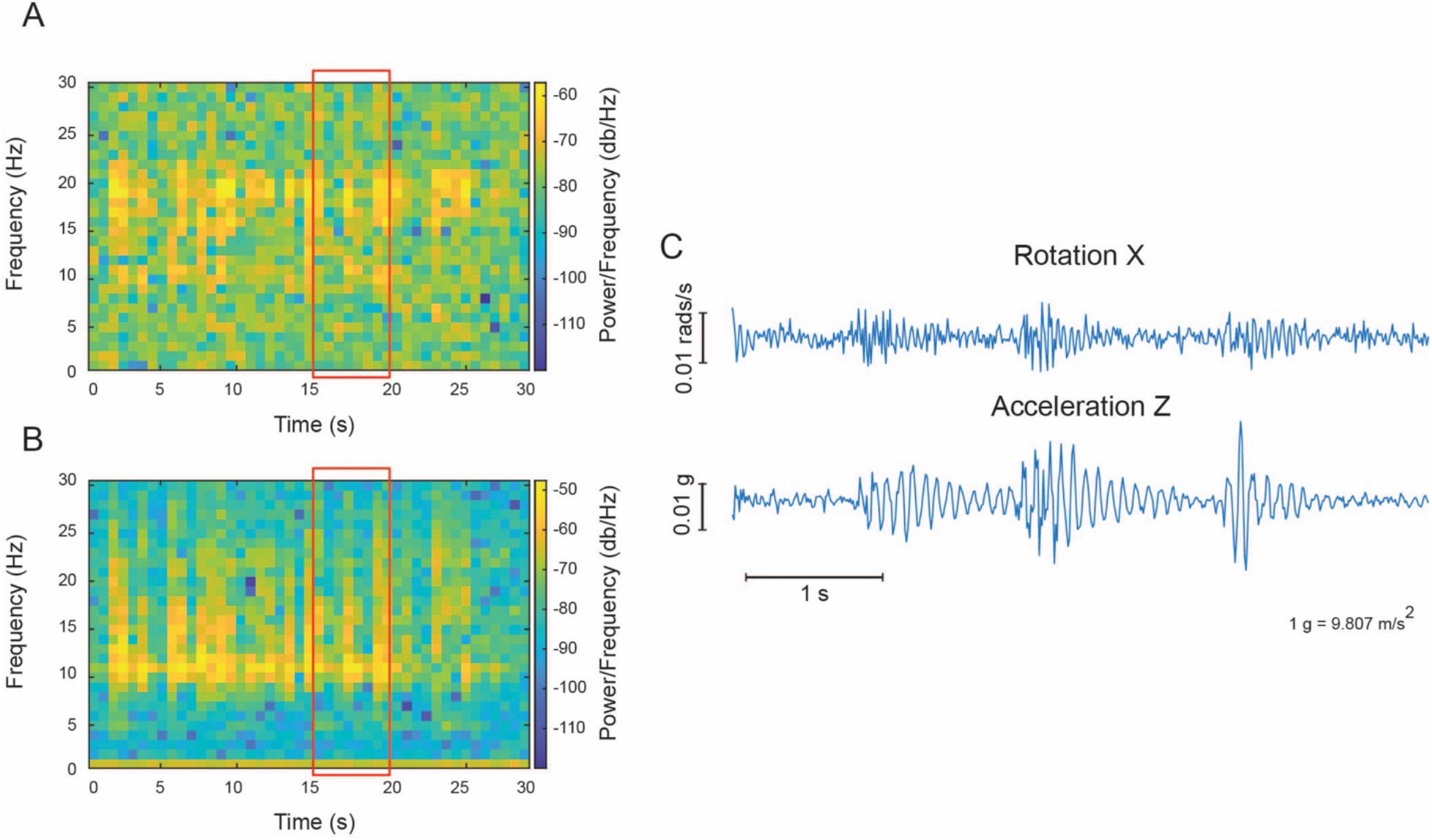
Simultaneous representative accelerometry data for variables ROTX and ACCZ, showing the quantified patterns in this study. **A.** Spectrogram for the accelerometer signal ROTX (rotation in the longitudinal axis of the cage). Notice the presence of circa 20 Hz rhythmicity for this rotation signal. **B.** Spectrogram for the accelerometer signal ACCZ (acceleration perpendicular the bottom of the cage). In this spectrogram, notice the presence of circa 10 Hz rhythmicity, appearing at similar times as rhythmicity in A. **C.** Data samples corresponding spectrogram region in the red rectangle. The samples show the relative timing of the ROTX and ACCZ oscillations, with three particular periods of enhanced oscillations, showing that the ROTX and ACCZ are recording the same mouse tremor phenomenon, yet with difference resonance conditions.

Simultaneous measurements of kinetics and kinematics were also performed to validate the kinetic data. As seen in Figure 3, the kinematic marker (tracking the top of the back) shows the same relative tremor frequency when compared to the frequency in accelerometry for ROTX (in this sample, a peak around 9 to 10 Hz for both signals). This confirmed that in our hands, the accelerometry data represented the animal’s tremor.

**Fig. 3.**
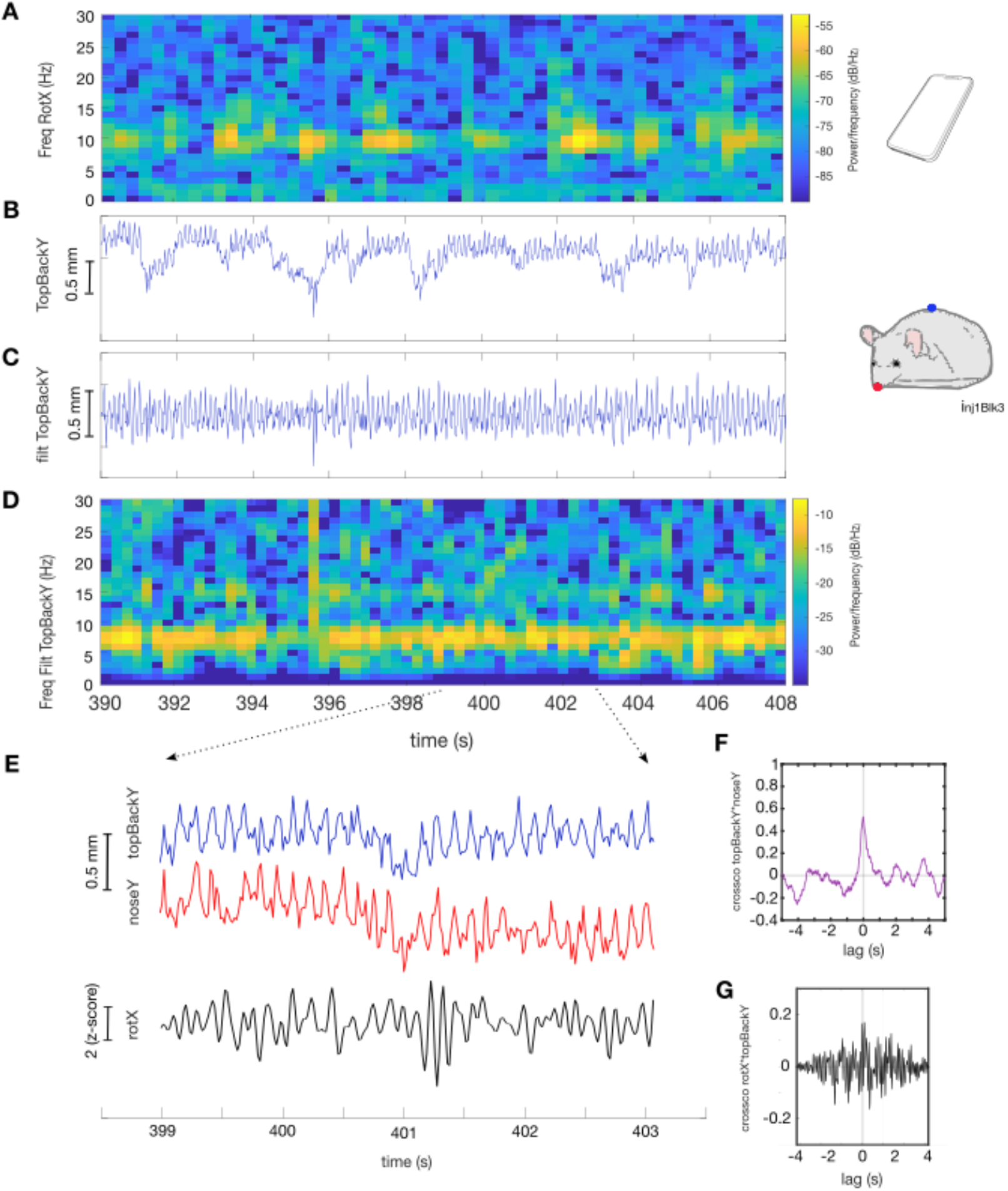
Simultaneous kinetic and kinematic tremor measurements in one mouse, showing the relative frequency match with each method. **A.** Spectrogram of the accelerometer-measured rotation in X (ROTX), which is along the length of the iPhone in line with the holding rubber-bands, via the Tremor12 application. **B.** Raw vertical position of the top-of-the-back label (TopBackY – the blue marker on the mouse sketch, adapted from a frame of the video), coming from DeepLabCut. **C.** Filtered (FIR equiripple high-pass > 3 Hz) vertical position of the same top-of-the-back (TopBackY) marker, to remove remaining slower displacements. **D.** Spectrogram of the filtered vertical position top-of-the-back marker in C, to emphasize the specific tremor frequency. **E.** Magnified portion (circa 5-s) of the time series for the simultaneous top-of-the-back (TopBackY - blue marker on the mouse in B, blue line) and nose marker (noseY - red marker on the mouse in B, red line) vertical positions. F. Cross-covariance between the top-of-the-back (topBackY) and nose (noseY) vertical positions for the time period in E (purple line), showing a largely in-phase (zero-lag) relation between the two body parts, at a given time.

After this confirmation for the dominant frequency bands in the pre-intervention data, we integrated between the 8-14 Hz and the 14-29 Hz frequency boundaries, averaged across all time windows per block. Results were initially focused on the second block of data between 5-10 min after harmaline injection, when we thought we would get the strongest tremor; afterwards we used the same methods on all blocks. Data for each mouse were then pooled into each group. A repeated-measures (pre vs. post-intervention) ANOVA with group (HIIT, RT, SED) as a factor served to determine the effects of the interventions on tremor.

#### Lactate and BDNF concentrations

Individual mouse concentrations for lactate and BDNF were compiled according to each group. An ANOVA with group as a factor was then carried out for lactate, as well as one for BDNF measurements.

## RESULTS

The three groups of mice received multiple tests and assessments before and after the intervention protocol (including the harmaline administration), to address the potential preventive effect of exercise on the amount of tremor in a mouse model of ET. We present descriptive data on the mice, the effect of the exercise protocol, and the effects of the interventions on tremor generated by the harmaline challenge. We also attempted to link the exercise effects with the brain physiology, via blood analysis.

### Descriptive data on the mice

We present data describing the groups of mice (individual age, weight, time on rotarod) in Table 1. Overall, the three groups of mice were of slightly different age [means (months) HIIT = 11.14; RT = 14.29; SED = 9.57; F(1,2) = 3.944, p = 0.038], but similar weight [means (g) HIIT =33.66; RT = 33.32; SED = 32.28; F(1,2) = 1.81, p = 0.192]. We did see an effect of the exercise protocol on the three groups on the weight [means (g) HIIT Pre = 34.44, vs post = 33.66; RT Pre = 33.57, vs. post = 33.32; SED Pre = 29.18, vs. post = 29.87; group by pre-post intervention interaction, F(1,2) = 14.689, p < 0.001]. This showed that exercise had an effect on the animals’ weight. In order to generally assess their balance and coordination, tests on the rotarod revealed no initial difference between groups [means (s) HIIT = 45.29; RT =26.71; SED = 31.57; F(1,2) = 1.172, p = 0.332], and no effect of the exercise protocol [means (s) HIIT Pre = 39.57, vs post = 45.29; RT Pre = 25.86, vs. post = 26.71; SED Pre = 29.14, vs. post = 31.57; group by pre-post interaction, [F(1,2) = 0.625, p = 0.547]. As such, no specific intervention showed an effect on rotarod performance. While the rotarod test was performed as usually indicated and at two speeds (31, 41), we must acknowledge that only one trial was taken pre- and post-intervention for each mouse.

**Table 1:** Descriptive data for the mice. The mean age for all mice was 12.33 months. Each group of mice (n=7) was weighed at the beginning of each week while in protocol. The descriptive statistics showed a decrease in the weight for the HIIT (means = 34.44 g vs 33.66 g) and RT (means = 33.57 g vs 33.32 g) groups while the SED showed an increase (means = 29.19 g vs. 29.87 g).

| Mouse | Group | Age at start of protocol (months) | Initial Weight (g) | Weight after protocol (g) | Rotarod @ 8 RPMs – (pre, s) | Rotarod @ 8 RPMs (post, s) | Rotarod @ 12 RPMs<br>(pre, s) | Rotarod @ 12 RPMs<br>(post, s) |
|---|---|---|---|---|---|---|---|---|
| 1 | HIIT | 11 | 39.71 | 38.32 | 109 | 116 | 113 | 120 |
| 2 | HIIT | 11 | 33.80 | 31.65 | 37 | 40 | 30 | 32 |
| 3 | HIIT | 11 | 30.25 | 29.68 | 48 | 45 | 50 | 53 |
| 4 | HIIT | 11 | 34.09 | 34.25 | 27 | 29 | 28 | 33 |
| 5 | HIIT | 11 | 41.3 | 41 | 27 | 31 | 25 | 27 |
| 6 | HIIT | 8 | 29.88 | 29.3 | 19 | 18 | 15 | 20 |
| 7 | HIIT | 15 | 32.05 | 31.42 | 26 | 29 | 26 | 32 |
| 8 | RT | 20 | 33.49 | 33.1 | 31 | 34 | 32 | 39 |
| 9 | RT | 7 | 32 | 31.78 | 12 | 12 | 10 | 13 |
| 10 | RT | 7 | 27.8 | 27.3 | 15 | 14 | 17 | 18 |
| 11 | RT | 20 | 39.9 | 39.88 | 26 | 26 | 25 | 23 |
| 12 | RT | 20 | 42.1 | 41.7 | 17 | 20 | 19 | 21 |
| 13 | RT | 20 | 29.2 | 29.4 | 37 | 39 | 34 | 33 |
| 14 | RT | 20 | 30.5 | 30.1 | 42 | 41 | 44 | 40 |
| 15 | SED | 7 | 28.9 | 29.6 | 29 | 28 | 30 | 21 |
| 16 | SED | 20 | 28.4 | 29.7 | 29 | 30 | 28 | 39 |
| 17 | SED | 8 | 27.8 | 28 | 31 | 33 | 37 | 38 |
| 18 | SED | 8 | 30.2 | 30.8 | 33 | 36 | 34 | 30 |
| 19 | SED | 8 | 29 | 29.3 | 12 | 10 | 13 | 10 |
| 20 | SED | 8 | 28.9 | 29.9 | 14 | 14 | 12 | 16 |
| 21 | SED | 8 | 31.1 | 31.8 | 39 | 40 | 50 | 67 |

### Effects of training on pre-post intervention exercise measurements

#### HIIT Group

To assess the potential training effects of this exercise intervention protocol, each mouse in the HIIT group underwent an exhaustion test before and after the intervention to assess changes in performance. Pre-training values ranged from 22 m/s to 28 m/s (mean = 24.3 m/s). After the 14-day HIIT protocol, the post-training values ranged from 24 m/s to 30 m/s (mean = 26.3 m/s). Data are in Fig. 4, showing that post-training speed was higher than pre-training speed (df = 6, t = -4.382, p = 0.005). This suggests that the HIIT training protocol permitted to increase the performance of the mice in this group.

**Fig. 4.**
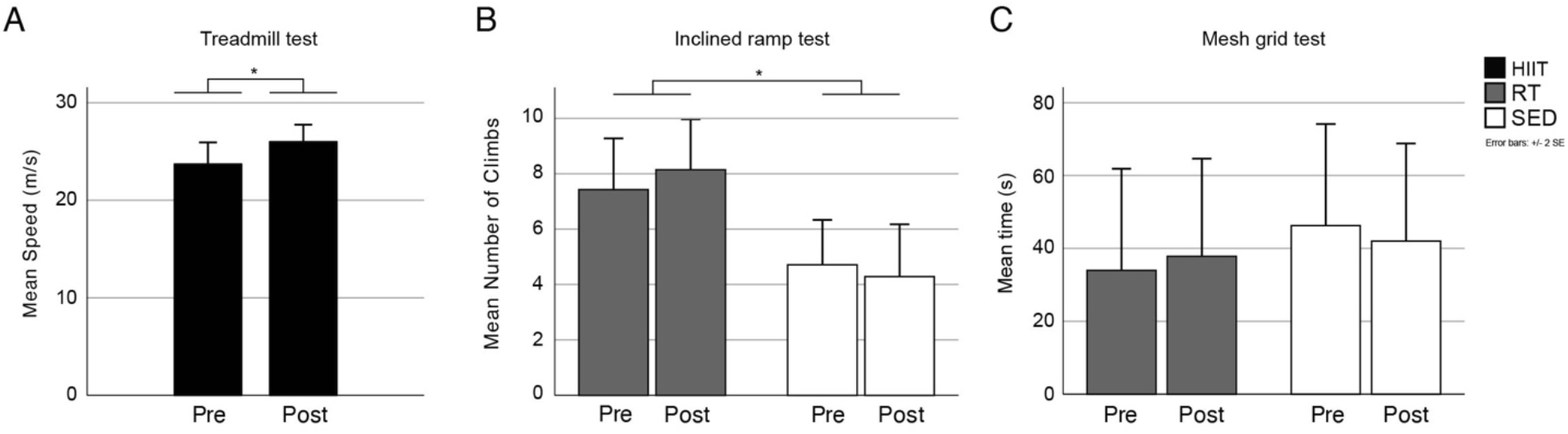
Testing (pre- and post-intervention) for each group. **A.** Maximal exhaustion testing for the HIIT group, before (Pre) and after (Post) undergoing the 2-week HIIT protocol. Bars represent the mean speed (m/s), with standard error, at which the mice reached exhaustion. The * indicates a difference < 0.05. B. and C. Inclined Ramp Mesh Grid and testing for RT and SED groups. **B.** Inclined ramp test (number of climbs) for the two groups, with no difference in the pre and post tests. There was a difference (* < 0.05) between the two groups. **C.** Mesh grid test (hanging time, in s) which did not show any difference between pre and post values, or between groups.

#### Resistance Training and Sedentary Groups

For the pre- post-intervention testing for the RT and SED groups, mice were tested on the inclined ramp (number of climbs) and inverted mesh grid (maximum time in that each mouse held onto the mesh, before reaching exhaustion) tasks. Data are shown in Figure 4, B & C.

#### Inclined Ramp

For the inclined ramp, the RT Group did not improve in the number of climbs after their training; in pre-intervention testing, the mice performed between 4-10 climbs while post-intervention, they showed a range of 5-12 climbs. The SED group also showed no improvement in climbs with pre-intervention ranging from 2-8 and post-intervention values remaining ranging between 1-8 climbs. However, the two groups (RT, SED) were statistically different [F(1,2) = 4.923, p = 0.047]; this is shown in Fig. 4B.

#### Mesh Grid

For the mesh grid (maximum time in s), the RT group held on for 0-90 s pre-intervention and 5-93 s post-intervention to the grid. The SED group held on a range of 13-120 s pre-intervention and 10-110 s post-intervention on the grid. The higher values of the SED were affected by the exceptional performance from Mouse 16. Data were not significantly different between groups [F(1,2) = 0.101, p = 0.756], as seen in Fig. 4C.

Overall, the pre-intervention and post-intervention data show a training effect for the HIIT treadmill group; while our data suggest that the RT group performed better in the RT testing, it appears their performance was not enhanced by the resistance training regimen. The SED group performance was lowest, and not affected by the passage of the intervention time period.

### Tremor assessment

#### Kinetics

Tremor was assessed by analyzing the variables of rotation in the cage longitudinal axis (aligned with the rubber bands supporting the cage, ROTX) and acceleration in the direction perpendicular to the cage bottom (ACCZ). As shown in Fig. 2, the signal had been first assessed to determine the specific boundaries which best represented the tremor power for the mice (ACCZ between 8-14 Hz – capturing the 10 Hz and ROTX between 14-29 Hz – capturing the 20 Hz). These represented simultaneous recordings of the same tremor (with different resonance because of the particular axis and variable sensibility). We tracked the changes across both variables and frequency bands, as the position of the animal in the cage could favor one variable over the other.

Since accelerometry and video data were recorded at the same time, the video data storage limitations guided how we separated the data, specifically in circa 5-min time blocks. We first analyzed block 2, which generally showed good power at our specific tremor boundaries. We considered this a useful first pass in assessing the effect of the harmaline injection; having tested these analyses on a block, we then analyzed the overall total of blocks available for each mouse.

#### Specific block analysis (block 2) Rotation (ROTX)

For ROTX, for the analysis of block 2, the repeated-measures (pre- and post-intervention) ANOVA with 3 groups (HIIT, RT & SED) and 2 tremor frequency bands (8-14 Hz, 14-29 Hz) revealed a significant pre-post effect [F(1,18)=4.546, p=0.047]. The power in the two bands was higher post-intervention than pre-intervention. As expected for ROTX, the main effect of frequency band was highly significant [F(1,18) =143.539, p<0.001] with a higher power in the 14-29 Hz band. The frequency band by group interaction was significant [F(2,18)=21.196, p<0.001] with the HIIT group showing the least difference of power between the two bands compared to RT and SED groups. A significant interaction was found between frequency band and the pre-post effect [F(1,18)=9.100, p=0.007]. Additionally, the pre-post by frequency band by group interaction was significant [F(2,18)=6.408, p=0.008]. Figure 5AB provides the description of this triple interaction. It is noticeable that the HIIT group showed a bigger difference across the two harmaline injections, with the 8-14 Hz band decreasing in power and the 14-29 Hz band increasing in power, when comparing with the RT & SED groups.

**Fig. 5.**
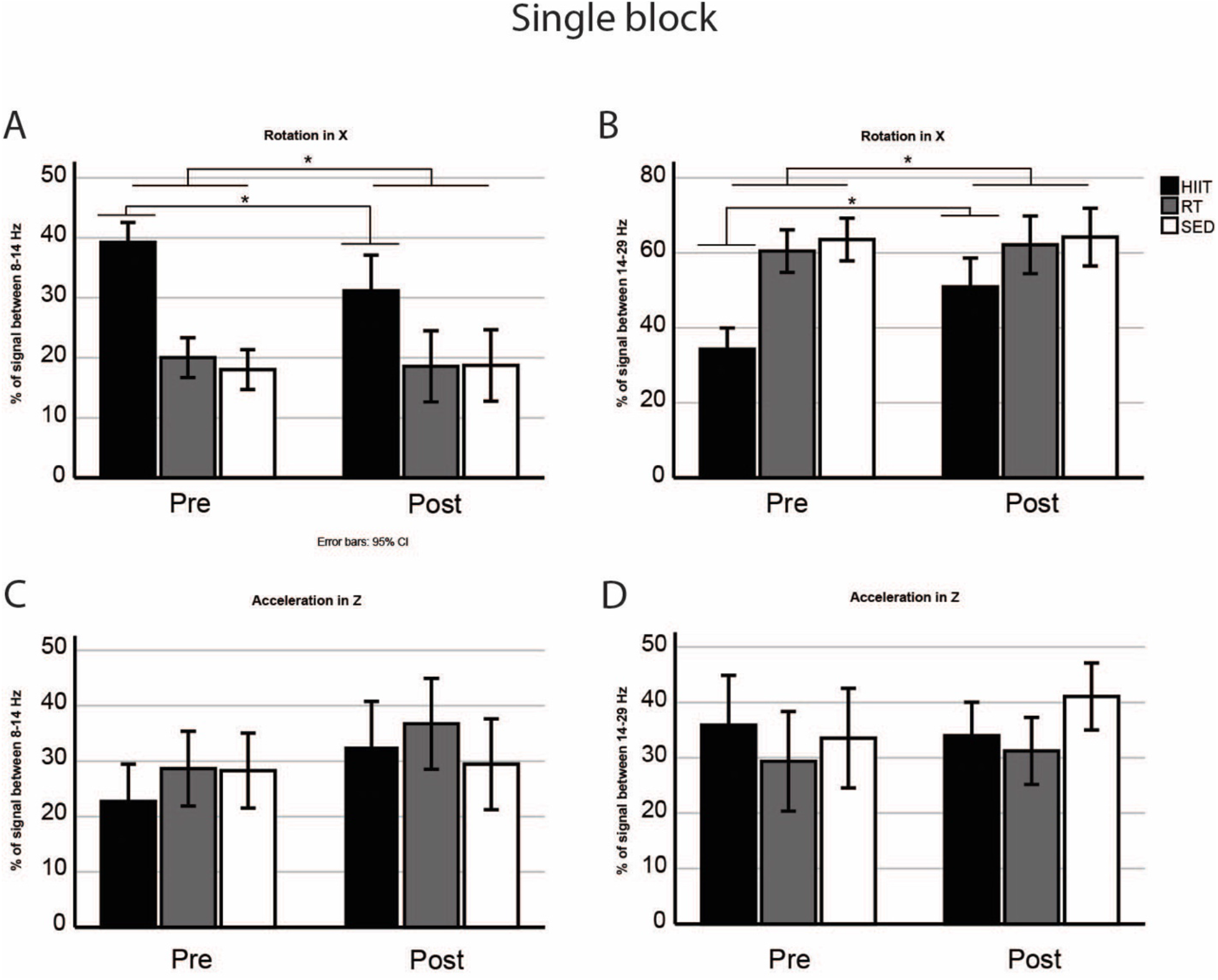
Tremor in a single block. Mean and Standard Error of power between 8-14 and 14-29 Hz in our three groups (HIIT, RT, SED), pre and post exercise protocols, using the ACCZ and ROTX variables, quantified in a single block (block 2). **A**: Power between 8-14 Hz of ROTX. **B**: Power between 14-29 Hz of ROTX. **C**: Power between 8-14 Hz of ACCZ. **D**: Power between 14-29 Hz of ACCZ.

To illustrate this specific change in the HIIT group, we are showing on Fig. 6 the spectrograms for one mouse, both from the ROTX values for the first block of data post-injection. Fig. 6A shows the accelerometer data before the intervention, while Fig. 6B shows the data post-intervention. The figure illustrates well an increase of about 5 Hz in tremor frequency for this mouse, after the exercise intervention.

**Fig. 6.**
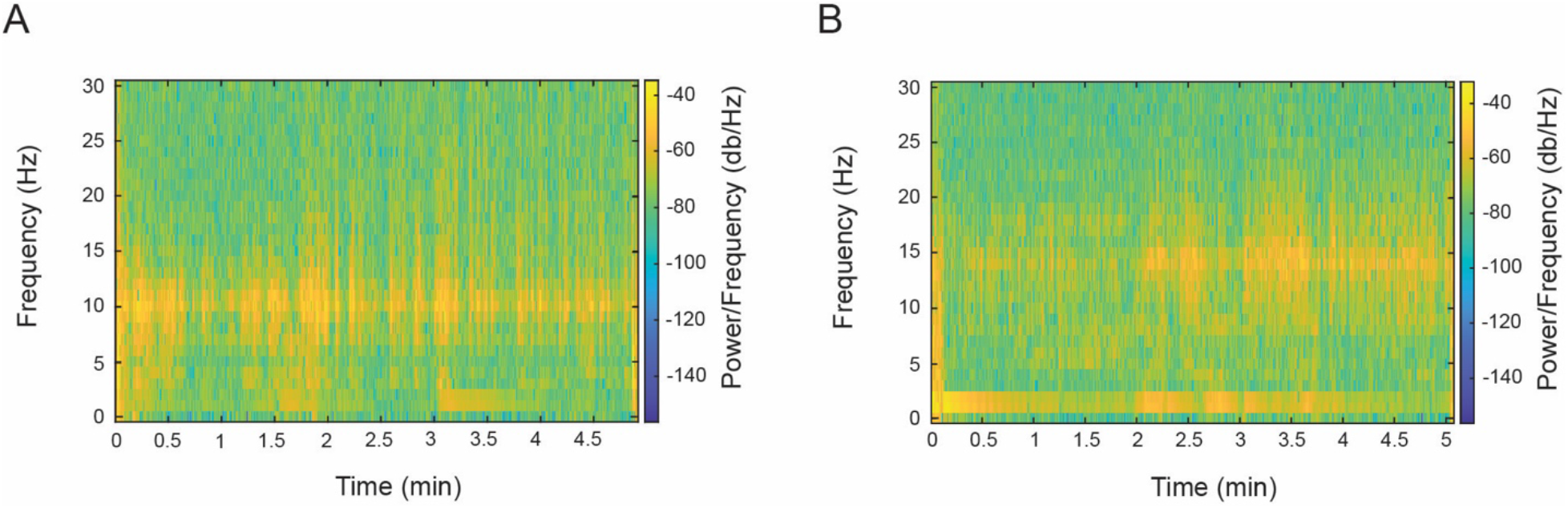
Tremor measurement (Rotation in X) for block 2 for one mouse in the HIIT group. **A**: Spectrogram for ROTX in the first post-injection period, before the exercise intervention. **B**: Spectrogram for ROTX in the first post-injection period, after the exercise intervention. This side-by-side view illustrates well the relative change in frequency band for the tremor, increasing frequency by about 5 Hz after the exercise intervention.

#### Acceleration (ACCZ)

For ACCZ, for the analysis of block 2, the repeated-measures (pre- and post-intervention) ANOVA with 3 groups (HIIT, RT & SED) and 2 frequency bands (8-14 Hz, 14-29 Hz) showed a significant main effect of group [F(1,18) =4.706, p=0.044], with the HIIT group overall yielding higher values as seen in Figure 5CD. No other significant difference could be discerned.

#### Overall Blocks Rotation (ROTX)

Analysis of the overall blocks also used a repeated-measures (pre- and post-intervention) ANOVA with 3 groups (HIIT, RT & SED) and 2 frequency bands (8-14 Hz, 14-29 Hz). Data are shown in Figure 7. For ROTX, there was a significant pre-post effect [F(1,18)=4.729, p=0.043] with a higher post-intervention power than the pre-intervention power, across the two bands. A pre-post by group interaction was also significant [F(2,18)=4.021, p=0.036]. In addition, the frequency band by group interaction was significant [F(2,18)= 23.667, p<0.001]. Lastly, the triple interaction (pre-post by frequency band by group) was also significant [F(2,18)=8.984, p=0.002]. To illustrate, Figure 7AB shows the notable difference between pre- and post-intervention in the HIIT group ROTX power between 8-14 Hz and 14-29 Hz: the integrated power between 8-14 Hz decreases post-training, while the 14-29 Hz integrated power increases.

**Fig. 7.**
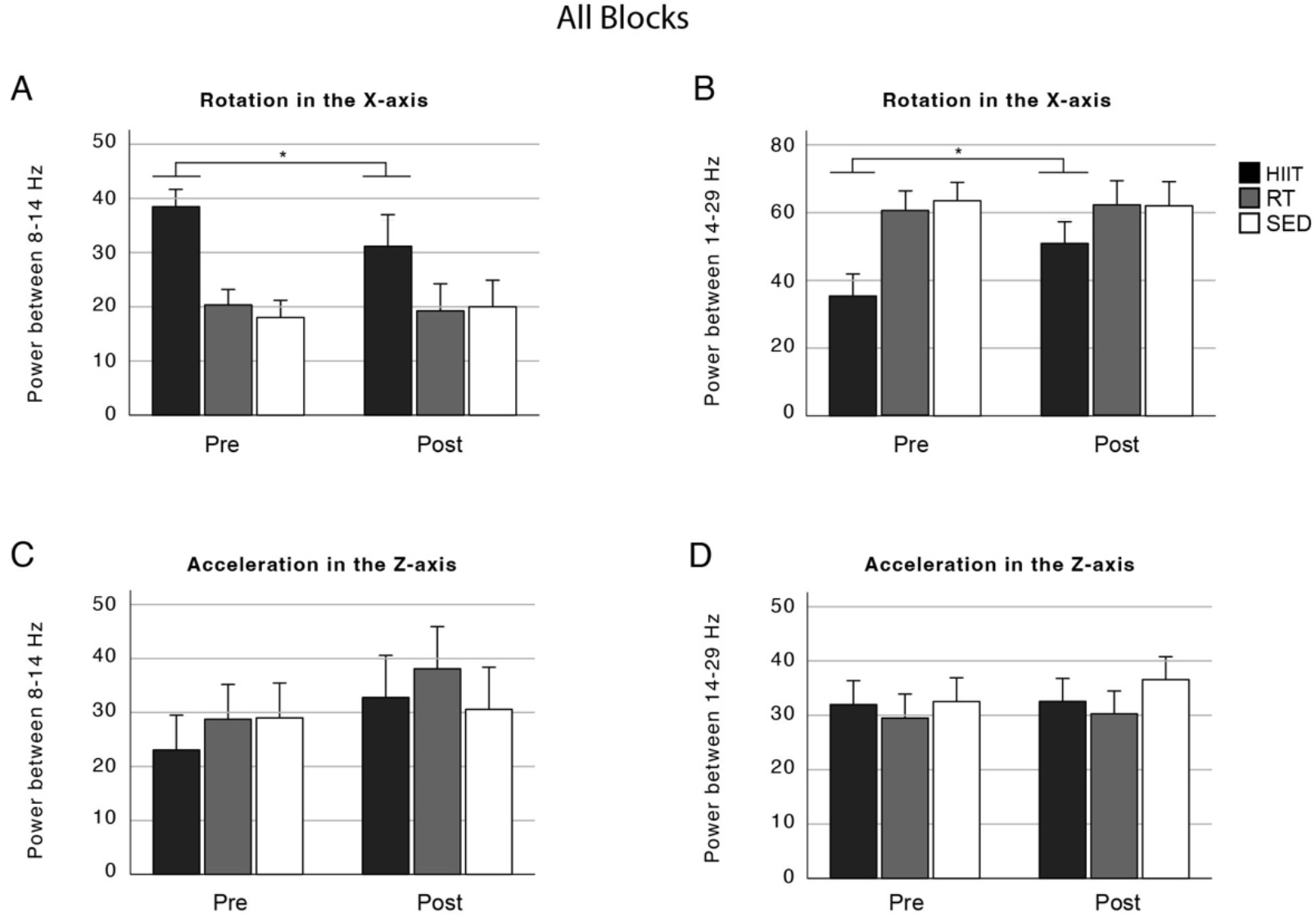
Tremor in all blocks. Mean and Standard Error of power between 8-14 and 14-29 Hz in our three groups (HIIT, RT, SED), pre and post exercise protocols, using the ACCZ and ROTX variables, averaged across all blocks. **A**: Power between 8-14 Hz of ROTX. **B**: Power between 14-29 Hz of ROTX. **C**: Power between 8-14 Hz of ACCZ. **D**: Power between 14-29 Hz of ACCZ.

#### Acceleration (ACCZ)

For ACCZ, the analysis showed a significant pre-post effect [F(1,18)=5.447, p=0.031], and a significant frequency band by group interaction [F(2,18)=3.66, p=0.046]. Other effects or interactions were not statistically significant.

### Lactate and BDNF

#### Lactate

For the lactate measurements (collected after the last exercise session for the HIIT and RT groups, and at a similar time for the SED group), we also ran a group comparison. The lactate concentrations were not statistically different between groups [F(1,2)=1.334, p=0.286], though we noticed a ranking of HIIT, RT and SED in order. Data are shown in Fig. 8A.

**Fig. 8.**
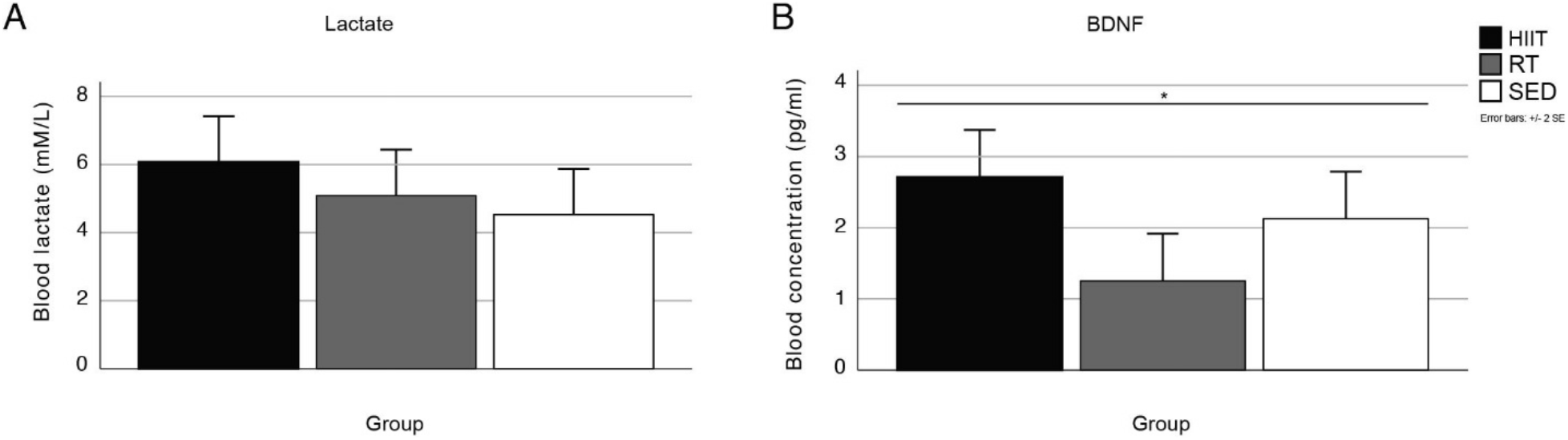
Lactate and BDNF measurements for all 3 groups (HIIT, RT, SED). These samples were collected at the end of the protocol. **A.** Mean & SE for lactate measurements. **B.** Mean & SE for BDNF measurements. * indicates a difference < 0.05.

#### BDNF

We also analyzed the BDNF levels from the blood samples collected from all mice in the 3 groups (HIIT, RT, SED), shown in Fig. 8B. From the ANOVA, the overall BDNF concentration was different between groups [F(1,2)=4.924, p=0.020], with the HIIT group showing the highest BDNF values.

## DISCUSSION

Exercise is increasingly considered as a useful preventive or curative component within someone’s lifestyle, and to support the individual with a neurological condition. In this precursor to a more circuit-specific enquiry in mice, and based on the harmaline model of essential tremor, we aimed to study the potential preventive effects of aerobic exercise in minimizing tremor. For translation ability, we used a titrated exercise approach (such as in the HIIT protocol), and found that the mice in the HIIT group showed exercise-related effects in tremor power across specific bands, which did not occur for the RT and SED groups. Indeed, when comparing pre- and post-intervention harmaline challenges, tremors in the HIIT group changed in frequency band relative predominance towards faster oscillations (i.e., for ROTX decreasing in the 8-14 Hz band and increasing in the 14-29 Hz band). From the literature, we were expecting mostly a decrease in power at the lower band (8-14 Hz): this change in band predominance was a surprise, yet this change could also represent a potential ‘neural resilience’ mechanism to decrease the impact of tremor in favor of the animal’s voluntary motor control. The following subsections will detail the measurable changes in performance brought about by the interventions, as well as the mechanisms underlying the effects of exercise on tremor.

### Effects of interventions on performance tests

To determine the efficacy of the interventions, we assessed the change in the adapted pre- and post-intervention testing. From the results, the HIIT exercise protocol was effective at improving the mice’s performance at the treadmill exhaustion test; however, we did not see a real improvement for the RT mice in their anaerobic performance (inclined ramp and mesh grid), despite our effort at making the mice work in force-generating tasks over a short time window, and with effort progression. We also saw that the SED group did not show a performance change at the resistance-related testing (also the inclined ramp and mesh grid). Thus, at least for the HIIT protocol, the exercise regimen was effective at improving the performance of mice: they likely benefited from the physiological advantages that this type of exercise can provide (42). In general, the physiological effects from increasing the multi-muscle force and whole-body metabolism coming from the HIIT exercise are likely distributed across the mice’s muscular and nervous systems. For the HIIT group, effects on the nervous system would likely come from a combination of metabolic and neural activation-specific changes, while for the RT intervention, we expected a greater emphasis on neural mechanisms (e.g., cerebrospinal and spinal) interactions (42) as changes to repeated exercise. The reason why the RT group did not improve could be related to the methodology we used, as we were unsuccessful in using weighted vests (43) onto our mice, instead opting for a time-constrained circuit approach to generate a progressive overload: this appears not to have been enough to trigger force improvements. The RT group was also our oldest group (there was a statistical age difference between groups), and their adaptability to exercise would have to be considered. However, aging did not seem to adversely affect the HIIT group, who were not the oldest, nor the youngest from our pool of mice.

### Effect of exercise on tremor

Our findings indicate that high-intensity interval exercise can attenuate the impact of tremor in the harmaline mouse model. Tremor kinetics showed a notable change for the group that underwent the HIIT protocol: the integrated power within 8-14 Hz decreasing and the 14-29 Hz power increasing. This can be interpreted as a band shift, as the intervention protocol modified the harmaline-induced tremor in its frequency spectrum, favoring higher bands for the HIIT group. From what we could see in the literature, this realignment is a first in the context of the effect of exercise on tremor. Considering the biomechanical relationships, a tremor frequency increase is usually an indication of an amplitude decrease (44) – which indeed has a positive effect on voluntary motor control.

The source of this change could come from a few different mechanisms, for example, from change to mechanical properties of muscles, to central generators, or to reflex feedback loops (45). More enquiries will be needed to parse out these options, as exercise can indeed exert influence at multiple levels. There is precedent for a change in frequency in ET patients, particularly in the context of aging effects: Elble (46) shows that frequency of tremor decreases with advancing age and duration from diagnosis onset. The corollary is that younger subjects have a higher frequency of tremor; as a potential mechanism, aging or ET-related changes could indeed affect the peripheral neural and muscular apparatus, limiting the ability to constrain tremor amplitude; the increase in amplitude then mechanically produces a decrease in tremor frequency. As we see a higher frequency band presence in the HIIT group after intervention, we can certainly purport that a potential mechanism for the influence of exercise could also be close to the effectors. Changes in the neuromuscular systems with the HIIT exercise could create a combination of muscle strength and better patterned activity in motor neural circuits, together improving motor control. Throughout the intervention, the HIIT animals could have thus gotten better at preventing larger amplitude tremors, with the effect of elevating the tremor frequency band. This could represent a preventive mechanism helping to support more fluid movement control, brought about by efficient exercise. The titration of preventive exercise should thus focus on reducing the oscillatory amplitude of tremor and increasing its frequency, in future studies.

### Potential neural mechanisms

Coherent with our own paradigm, the HIIT exercise could also have modulated specific sensorimotor circuits that generate the tremor. The harmaline model of tremor strongly activates olivocerebellar connections (9). Olivocerebellar afferents are modulated by cerebellar nucleo-olivary pathways (47), in turn under inhibitory control from cerebellar cortex Purkinje cells. In a modelling framework, disturbing GABAergic receptors in the dentate nucleus of the cerebellum successfully adjusts the tremor frequency. Indeed, Zhang and Santaniello (48) show that amplitudes and decay times of GABAergic currents to the dentate nucleus can promote sustained oscillatory activity at tremor-related frequencies. Thus, a dysfunction of inhibitory signaling in the cerebellum can contribute to the generation of rhythmic tremor activity. Exercise-related neuroplasticity could also influence GABAergic inhibition in specific circuits. According to Maejima et al. (49), aerobic exercise increases BDNF expression in the hippocampus, modulating GABAergic synapses, with the net effect of contributing to hippocampal plasticity.

According to Yu and Krook-Magnuson (50), the connectivity of the hippocampus and cerebellum may be a possible connection to the neuroplasticity effect of BDNF. One implication is that through neural and/or exerkine modulation, the HIIT protocol could help adjust GABAergic function, potentially in the cerebellum, thereby adjusting the muscular oscillatory output and tremor. This adjustment of inhibition at the local circuit level would adjust tremor-frequency reverberation across the cerebello-thalamic networks.

Inspecting the circuit interactions, the attenuation of tremor in the harmaline model could be explained by the modulation of the IO cell output through a nucleo-olivary GABAergic inhibition (47). Pan et al. (51) discussed the modulation of climbing fibers to Purkinje cell activity leading to tremor generation: excessive cerebellar oscillations (4-12 Hz) from olivocerebellar inputs would be a key feature of tremor generation in ET. The HIIT protocol may also contribute to the reduction of hyperactivity in the pathway by synchronizing neural firing in a way that prevents the pathological oscillations responsible for tremor. A mechanism by which this could happen is by adjusting the activity of climbing fiber synapses, as climbing fiber synaptic pruning deficits were implicated in the excessive cerebellar oscillations. While in our case, a two-time harmaline administration might not have triggered a massive synaptic reconfiguration as seen in mutant mice, the neuroplasticity from the intense voluntary controlled exercise in the HIIT protocol could potentially adjust synapses to prevent the reverberation. This exercise-related synaptic plasticity could be through the presence of circulating BDNF across the cerebellar circuits, combined with repetitive activation through the execution of treadmill running.

BDNF is characterized as an exerkine—a bioactive substance synthesized during exercise implicated in neuroprotection (27). It is important to note that even though BDNF is not believed to cross the blood-brain barrier, other exerkines such as lactate act through BDNF-mediated mechanism in the brain (28). Other BDNF mechanisms promoting neuroprotection can be involved, such as the NMDAR/Ca2^+^ cascade, where BDNF works as a regulator to reduce toxic NMDA receptor signaling triggering a neuroprotective effect (52). While it is only one of the many exerkines, we did find a difference in blood BDNF concentration after our exercise protocol. Of note, BDNF activates neuronal differentiation from stem cells and impacts mitochondrial bioenergetics in the brain: its action activates the peroxisome proliferator-activated receptor γ coactivator-1α (PGC-1α), which increases mitochondrial biogenesis and ATP and NAD+(53). In addition, lactate is involved in mitochondrial bioenergetics through the activation of SIRT1 and engaging the PGC-1α pathway. The HIIT exercise intervention (specifically, and not the RT protocol) could have upregulated PGC-1α, which is a crucial pathway for mitochondrial biogenesis, energy metabolism and the enhancement of neuronal resilience. The increase of activity in this pathway would promote the production of lactate, which crosses the blood-brain barrier to stimulate BDNF expression in the regions controlling tremor – here in many neurons across the sensorimotor networks, but a first enquiry for us could be through cerebellar nucleo-olivary pathways. As we did not find a change in lactate concentrations, an adjustment in the protocol for lactate collection might be in order: it could very well be that we collected lactate too late after the exercise to capture its full effect.

### Limitations

There are several limitations of this study. First, we indeed have a sample of mice that was biased towards females (F: n= 19, M: n=2). This imbalance excludes a direct comparison between sexes. An equal number would need to be considered in future studies to look at the effect from potential factors such female hormones (e.g., progesterone). The response of harmaline in the context of higher female or male hormones would be valuable for the gap in the literature. The effect of sex hormones on neurotransmitters such as GABA, serotonin, dopamine and glutamate has been studied, but there is scarcity of studies on harmaline effects (54). Other limitations included the timing of the lactate measurement relative to exercise, which is a more time-sensitive metabolic indicator, when compared with BDNF, whose quantity can be assessed over days after the training. Lactate was not taken right after each respective post-intervention exercise protocol, since our design mandated a cool-down period after this session; therefore, at minimum a 20-min gap occurred between end of exercise and the tail-bleed. It would also have been ideal to take a relative lactate measurement, comparing before and after the exercise, to track a change in lactate expression. Lezi et al. (55) injected lactate IP before and after an exercise protocol – with lactate mediating changes in brain bioenergetic-associated parameters. Future studies can focus on optimizing these particular blood biomarkers, on the particular circuits affected by high-intensity whole-body exercise to attenuate tremor, and even consider the study of BDNF receptors at particular cerebellar synapses, such as in Purkinje cells (56) or through its effects excitatory-inhibitory circuit differentiation (57).

## CONCLUSION

This study investigated the value of exercise (and HIIT) as a preventive measure for essential tremor, with a focus on the harmaline mouse model. We report a change in the power of tremor across two frequency bands from the HIIT exercise protocol, likely modulating the activation of the tremor network, in favor of voluntary motor control. Exercise could thus influence the expression of tremor, as we saw in our mice: such an intervention has great potential for translating to human studies. Further in-depth physiological studies would need to be conducted to find the network hubs more responsible for this resilience: in addition to improving the musculoskeletal systems, exercise could help coordinate the central generators and the interconnected reflex pathways. We focused on the central olivo-cerebellar pathway, as we injected harmaline in mice. Future investigations to explore other genetic models of ET (51), in the structured contribution of aging and tremor, as well as in addressing exerkine signaling across networks, would help clarify the physiological effects of exercise in ET. Optimized exercise could become a valuable adjunctive treatment for individuals with ET and/or other motor disorders, paving the way for non-pharmacological, accessible and active options to complement other interventions.

## ACKNOWLEDGMENTS

We thank Joseph Gerges and Michael Lagendyk for help in data acquisition and analysis. A special thanks to Amanda Brown, cAHT, for her help and guidance in animal care and technical support. Thanks as well to Marie-Ève Rivard (Manager, Clinical Analysis Suite) and the School of Health staff for consultation and work on the blood sample analyses. Thank you to Eva Carlsen for courteous advice on the kinetics measurements.

## AUTHOR CONTRIBUTIONS

RC and AB conceived and designed research; SC and RC performed experiments; SC and RC analyzed data; SC, RC and AB interpreted results of experiments; SC and RC prepared figures; SC and RC drafted manuscript; SC, AB and RC edited and revised manuscript; SC, AB and RC approved final version of manuscript.

